# Experimental analysis and biaxial biomechanical behaviour of ex-vivo sheep trachea

**DOI:** 10.1101/2021.11.26.470180

**Authors:** Fulufhelo Nemavhola, Harry Ngwangwa, Thanyani Pandelani

## Abstract

**Purpose:** The purpose of this study is to investigate the mechanical behaviour of the tracheal tissue under biaxial tensile loading. Furthermore, the study examines the material properties of the tissue through a study of the model parameters for six constitutive models.

**Materials and methods:** The fourteen (n = 13) trachea sheep (Vleis Merino) pieces of tissues measured to be ^~^ 30 × 20 mm where only the effective area subjected to engineering strain was ^~^ 25 × 16 mm. In this study, we assume that the tracheal tissue is anisotropic and incompressible, therefore we apply and study the material parameters from six models namely the Fung, Choi-Vito, Holzapfel (2000), Holzapfel (2005), Polynomial (Anisotropic) and Four-Fiber Family models.

**Results:** The results show that the trachea tissue is twice as stiff along the circumferential direction as it is along the longitudinal direction. It is also observed that the material properties are different (non-homogeneous) along the trachea.

**Conclusions:** The findings of this study will benefit computational models for the study of tracheal diseases or injuries. Furthermore, these findings will assist in the development of regenerative medicine for different tracheal pathologies and in the bioengineering of replacement tissue in cases of damage.

## 1. Introduction

The normal physiological functioning of the trachea has led to more research focussing on the understanding of its mechanical behaviour under normal airflow conditions. As a result, most research has focussed on the study of fluid-structure interaction between the air and the tracheal tissue [1–6]. These studies have examined and analysed how the tracheal wall interact with the air as it gets pulled into and pushed out of the lungs. In such studies the focus has been on understanding tracheal flow by employing computational fluid dynamics methods. However there remains questions on what part of the normal breathing action does the tracheal wall really play or if a damaged but open tracheal wall still perform its function. When one feels the trachea during deep inhaling and exhaling exercise, the tracheal rings can be sensed moving up and down along the neck which implies that the tracheal muscle plays a part during inhaling and exhaling processes. Such subtle actions of the trachea that make breathing feel normal to an animal are a focus area in the development of replacement material for the tracheal tissue. The understanding of the working mechanisms of native trachea will elucidate the development of the tissue-engineered trachea which remains unclear [7], however relevant mechanical and biomechanical properties of the trachea are incompletely characterised [8].

To understand fully the mechanical properties of the soft tissues, it is necessary to understand its mechanical behaviour [9–12]. Mechanical properties of soft tissues may be utilised in developing detailed computational models to study mechanisms of diseases [13–18]. But how the implantation of a replacement material affects the response of the physiological response of a trachea is a challenge. In this study, the mechanical behaviour of the tracheal tissue under biaxial tensile loading is investigated. The stress-strain behaviour of the tissue in the circumferential and longitudinal directions are examined and discussed. Due to its very low physiological loading, often below the limit stresses in the toe region, the strains are kept to within 30 % strain.

Besides studying the stress-strain behaviour of the tracheal muscle, the mechanical properties of the tissue are studied by examination of the material parameters and performances of six different constitutive material models. The Fung, Choi-Vito, Holzapfel (2000), Holzapfel (2005), Polynomial (Anisotropic) and Four-Fiber Family hyperelastic constitutive models have been previously utilised in studying mechanical behaviour of the soft tissues [19, 20]. The material parameters obtained from the selected hyperelatic constitutive models are normally utilised in developing finite element models [16, 21–25]. It is expected that the analysis of the trachea using constitutive models may improve surgical outcomes. Within the limitations of such a study, there are two very important findings: firstly, the tracheal muscle is stiffer along its circumferential directions probably due to the circular cartilaginous rings [8, 26–28] which are more rigid than the adjoining soft tissue around it; secondly, the material is highly random and non-homogeneous as exhibited by the wide margin in the standard deviation of its material parameters across the different test specimens. In drawing the latter conclusion, the authors are mindful of the fact that there could be other factors that may have influenced such a finding although the stringent control conditions under which the testing was carried out may have eliminated such eventualities. The tested tissues showed anisotropy along the circumferential direction at larger strains which is typical for soft tissues [29]. Although it might be expected that the same trend would also be true for the longitudinal direction, it is hard to state that in the present study since it seems that 30 % strain was not large enough to stretch the tissue along the longitudinal direction beyond its toe region.

The study is aimed at investigating the mechanical behaviour of the windpipe (trachea) in two directions: aligned along its circumference and in the direction of the airflow. The trachea serves two important purposes primarily as an air conduit into and out of the lungs [5] and secondly as a preconditioner and prefilter for inhaled air. In its normal physiological functioning, the trachea is not subjected to excessive mechanical loads in any of these directions. It is rather subjected to very low gauge pressure which may cause stresses well below the limit of its toe region, which were calculated as 3.3 kPa in tension during inspiration, −5.25 kPa in compression during expiration and 7.2 kPa for mechanical ventilation [30]. However, there might be several traumatic situations and illnesses that subject the tracheal tissue to very high stresses in the circumferential, longitudinal, or radial directions. A comprehensive literature study of the published cases of tracheal replacement between 1898 and 2018 on 290 patients reported that a breakdown of large tracheal defects comprised cancerous tumour invasions (60.4 %), airway trauma (6.7 %), critical airway stenosis (7.8 %), congenital stenosis (6.0 %), tuberculosis (3.9 %), prolonged intubation (3.9 %), and other conditions (11.3 %) [5, 31]. The most common tracheal injury (tracheal stenosis) [32] is usually caused by a tracheotomy intubation with unsuitable pressure. Despite the availability of various treatment procedures, with varying degrees of success, it is reported that even after treatment, the stenosis may reappear especially in cases of serious pathologies [33–35]. Such problematic situations with respect to the total treatment of tracheal injuries makes it necessary to understand its mechanical behaviour.

## 2.0 Materials and methods

### 2.1 Tissue acquisition and preparation for mechanical testing

The trachea was delivered to University of South Africa (Unisa) Biomechanics Laboratory after the slaughtering of sheep (Vleis Merino) of between 40 000 and 42 000 grams. The delivery of fully sheep trachea was delivered within 2 hours of slaughter from the abattoir in a temperature-controlled bag. The temperature was kept at maximum of 5°C during transportation. The fourteen (n = 13) trachea pieces of tissues measured to be ^~^ 30 × 20 mm where only the effective area subjected to engineering strain was ^~^ 25 × 16 mm (See Figure 1). In this study, the microstructural coordinates were not utilised because there was no imaging to ascertain the exact fiber direction. Therefore, two directions were defined as per the coordinate system of the trachea where longitudinal direction was along the length of the trachea and the circumferential direction was defined to be around the circumference of the trachea.

**Figure 1.**
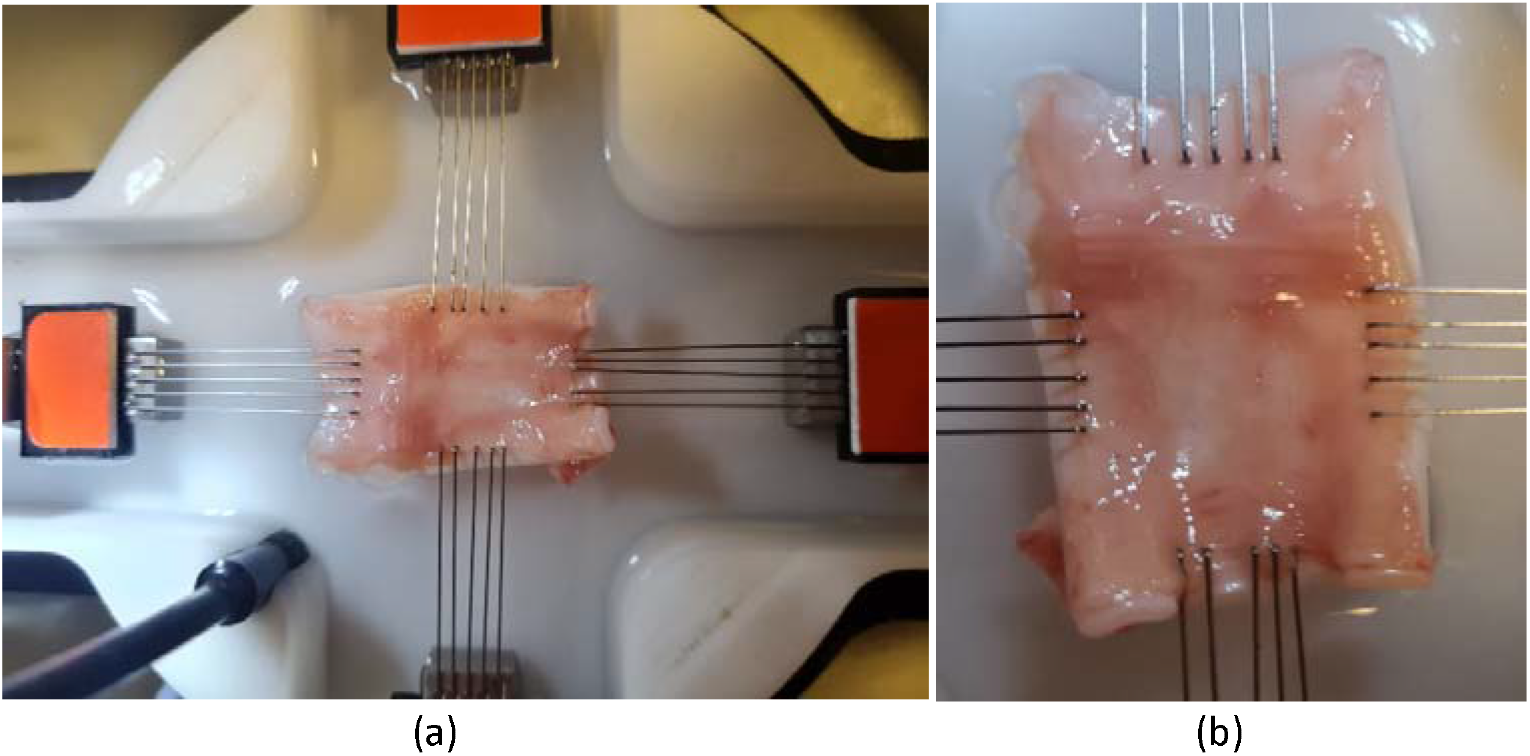
Experimental set-up of trachea subjected to biaxial tensile loading. (a) The 30 × 20 mm sheep trachea cut tissue hooked by the CellScale Biaxial testing system (BioTester 5000 CellScalle, Wateroo, ON, Canada^®^) submerged in the with Saline 0.91% w/v of NaCl was heated to 33°C before the sample tissue is placed in the bath for mechanical testing (b) magnified sheep trachea cut tissue hooked at ^~^ 26 × 15 mm.

### 2.2 Biaxial mechanical testing

To capture the mechanical properties of the sheep trachea, the CellScale Biaxial testing system (BioTester 5000 CellScalle, Wateroo, ON, Canada^®^) was utilised by applying eqi-biaxial forces on the excised trachea tissue. In this study, the microstructural coordinates were not utilised because there was no imaging to ascertain the exact fiber direction. Therefore, two directions were defined as per the coordinate system of the trachea where longitudinal direction was along the length of the trachea and the circumferential direction was defined to be around the circumference of the trachea. The excised tissue was cut in approximately 30 mm and 20 mm in longitudinal and circumferential directions, respectively. These tissues were placed in the biaxial device that has hooks attached to a pully system as reported previously [20, 36, 37]. Vernier calliper was utilised in capturing and measuring the thickness of the tissue in four different points where the average thickness was then utilised for further processing of engineering stresses and strains. As previously reported, in order improve accuracy of stress and strain data, the preloading of 10 % at the strain rate 0f 0.001/seconds. Saline 0.91 % w/v of NaCl was heated to 33°C before the sample tissue is placed in the bath for mechanical testing [19, 38, 39]. A 0.005 N equi-biaxial preload was applied to both longitudinal and circumferential directions. Finally, the strain rate of 30 % strain/10 seconds qui-biaxial loading and recovery was applied on the tissue.

### 2.3 Biaxial Mechanical testing

## 3. Theoretical formulation

### 3.1 Tissue stress-strain analysis

In this study the stresses were calculated through the first Piola-Kirchoff stress T in the two-directions using the equation:

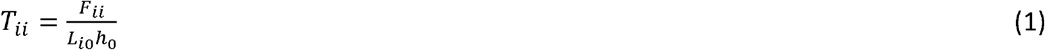

Where F is the load vector in directions i=1,2 and for the current study these indices represent longitudinal (cross-fibre-direction) and circumferential (fibre-direction) direction. The index 0 denotes the undeformed state. L represents tissue length, and h represents tissue thickness.

The finite strains were calculated by the formula:

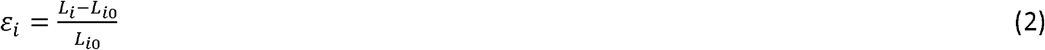

The calculated stress results are cut-off at 30% strain. These stress results however are noisy therefore they were further filtered with an 8-point moving average filter in Excel. The data were resampled and further smoothened using a quadratic function.

### 3.2 Constitutive modelling

The passive response of a biological soft tissue is more complex than the response of elastic solids due to the fact that they undergo finite deformations under mechanical loads [29, 40]. Thus, the passive phases of the tissues’ deformations are considered using the nonlinear theory of hyperelasticity. The most useful quantity in deriving the expressions for the passive response of materials that undergo finite deformation is the strain energy function [29]. There is a huge number of variations of its implementation in the literature depending on different cases. In this study, we assume that the tracheal tissue is anisotropic and incompressible, therefore we apply and study the material parameters from six models namely the Fung, Choi-Vito, Holzapfel (2000), Holzapfel (2005), Polynomial (Anisotropic) and Four-Fiber Family models. In the following sections, these models are presented in terms of their strain energy functions.

#### 3.2.1 The Fung model

The Fung model is a hyperelastic anisotropic material model for stress-strain description of arterial wall. It is phenomenological in nature and its incompressible form is given by [40, 41].

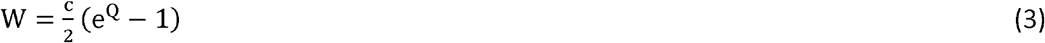

Where 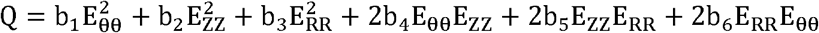; and b_i_ are the material parameters.

#### 3.2.2 Choi-Vito model

Choi-Vito model is hyperelastic anisotropic material model developed for canine pericardium. The model is fully phenomenological and formulated through the components of Green-Lagrange strain tensor. The model is implemented in an exponential format and its strain-energy function is expressed as [42].

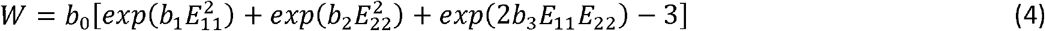

Where *b_i_* are the material parameters.

#### 3.2.3 The Holzapfel (2000) model

This model is hyperelastic anisotropic material model for stress-strain description of arterial layers. It is constituted by forms of the strain energy function that represent isotropy and anisotropy. Thus, its strain energy function is given by [43].

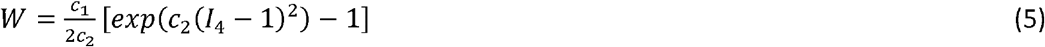

Where *c_i_* are the material parameters. The model is implemented in an exponential format.

#### 3.2.4 The Holzapfel (2005) model

This model is a variation of the previous model also aimed at modelling stress-strain behaviour of hyperelastic anisotropic materials, specifically the arterial layers. This model models the arterial walls as composite material with spirally arranged layers. Its strain energy function is expressed as [44]

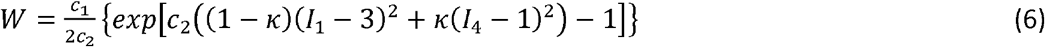

Where *c_i_* are the material parameters and *κ* is a parameter that modulates the convergence rate.

#### 3.2.5 The four-fiber family model

This model is hyperelastic anisotropic material model for stress-strain description of aortas and aneurysms. The model represents an elastin-dominated amorphous matrix reinforced by four families of (collagen) fibers (in axial, circumferential and diagonal directions) whose strain energy function is expressed as [45, 46]

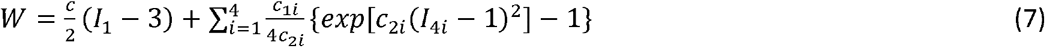

This model implements a hybrid polynomial and exponential format where *c*, *c*_1*i*_, *c*_2*i*_ are material parameters.

#### 3.2.6 The polynomial (anisotropic) model

Like its name, this model is hyperelastic anisotropic material model whose strain energy function is expressed as a polynomial series of isotropic and anisotropic strain invariants given as [47].

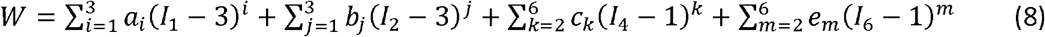

Where *α_i_*, *b_j_, c_k_*, and *e_m_* are material parameters.

### 3.3 Data analysis

In this study, we use the constrained optimisation by linear approximation algorithm (COBYLA (3^rd^ party: SciPy)) implemented in Hyperfit software for fitting the equi-biaxial tensile experimental data of Fung, Choi-Vito, Holzapfel (2000), Holzapfel (2005), Polynomial (Anisotropic) and Four-Fiber Family hyperelastic constitutive models given in Equations (3)–(8). Five metrics that measure models’ performances are evaluated for each model. These metrics are coefficient of determination, correlation coefficient, Normalized Root Mean Square error, Normalised error, and evaluation index. Their expressions are given in Equations (9)–(14) below.

The coefficient of determination (R^2^) (also known as Nash-Sutcliffe coefficient) is defined as follows:

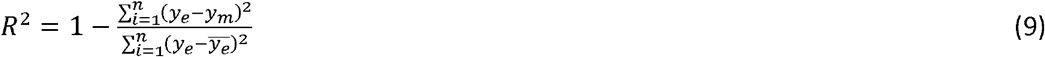

Where *y_e_* is the experimental data, *y_m_* is the model predicted data, 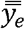 is the average value of the experimental data, the indices *i*, …, *n* denote the data points, and *R*^2^*ϵ*〈−∞, 1〉, where a perfect fit is defined for *R*^2^ = 1.

The Evaluation Index (EI) is a critical parameter in evaluating how the hyperelastic constitutive model fits the experimental data, and is given by

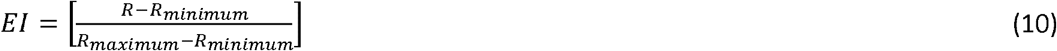

where,

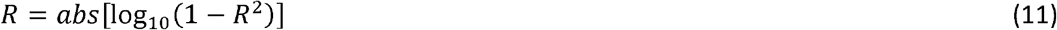

The *R_minimum_ and R_maximum_* represent the *R* values for poorest and best fitting hyperelastic models, respectively. EI in Equation (11) is a comparative parameter whose values may span values between 0.0 for poorest fitting models, and 1.0 for best fitting models. Therefore, the higher the coefficient of determination (R^2^), the higher the model fit (EI).

The correlation coefficient (r) is defined as

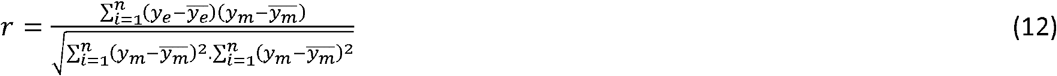

Where 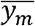 is an average value of the model predicted data, and all the other quantities in Equation (12) are defined as given in Equation (9).

The Normalized Root Mean Square error (NRMSE) is defined as

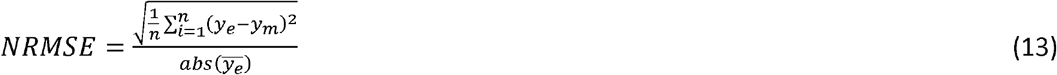

The Normalised error (NE) is expressed as

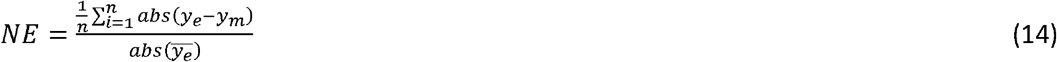

## 4. Results

Stress-strain curves for the trachea muscle subjected to equi-biaxial tensile loading are plotted in Figure 1. The stress-strain curves for trachea muscle in the figure show that the circumferential direction has a well-defined nonlinear behaviour as the tissue gets stretched from the toe region to 30% strain. At 30% strain, the tissue in the longitudinal direction still appears to be within the toe region with the stress-strain curve showing much more tissue compliance. The tracheal tissue in the longitudinal direction attains 30% strain below 50 kPa while the same level of strain is attained in the circumferential direction at stresses of about 100 kPa (double as much as that in the longitudinal direction). These findings are in line with previous studies by other researchers on the relatively higher stiffness of the tracheal tissue along the circumferential direction [8, 48] as compared to the longitudinal direction. Along the circumferential direction the toe region is only up to 15% strain as opposed to the 30% strain for the circumferential direction. In the previous studies it was found that the cartilaginous rings were stiffer towards the cranial regions than towards the caudal regions [27].

**Figure 1.**
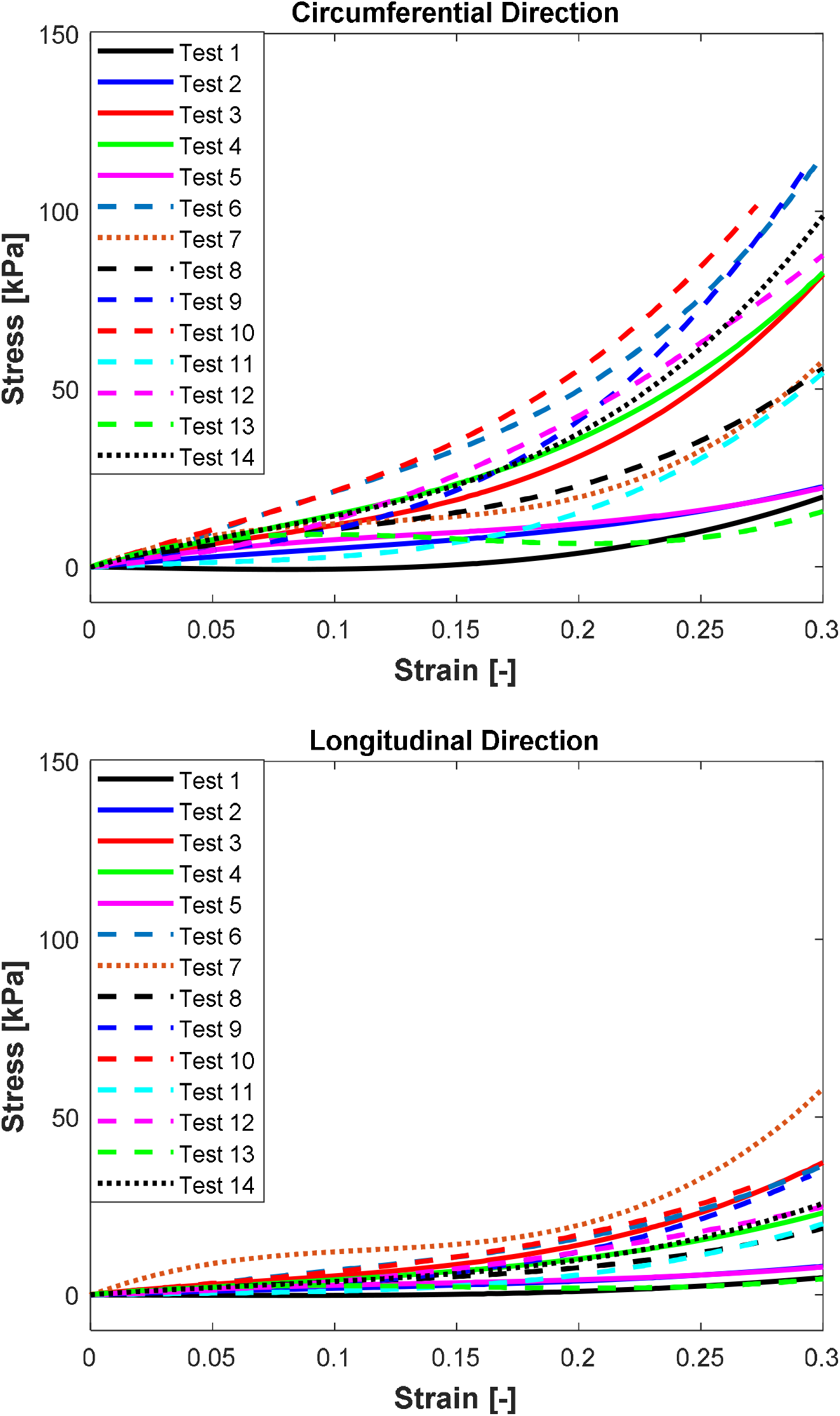
N =13 sheep trachea subjected to equi-biaxial mechanical forces shoing stress vs strain mechanical properties at 30 % strain (a) Circumferential and (b) Longitudinal. Test 7 showing the highest stress and Test 1 showing the lowest stress in the longitudinal direction, respectively. Additionally, test 10 and Test 1 showing highest and lowest tress in the circumferential direction, respectively.

In terms of anisotropy, it is extremely difficult to establish from this test the anisotropy of the tissue along the longitudinal direction since it appears the tissue is still in its toe region. But along the circumferential direction, the tracheal tissue show anisotropy. The tissue anisotropy grows with strain level in both directions although the circumferential direction is clearly more anisotropic from strain ranges greater than 15% strain. It is not clear what might have caused the extraneous stress-strain bevaviour of the specimen number 7 along the longitudinal direction, although its behaviour along the circumferential direction falls within the ranges of all the other specimen stress-strain curves.

Figures 2–4 plot the average values of the coefficient of determination (R^2^), the normalised root mean square error (NRMSE), and the evaluation index that were calculated over all the fourteen tests for each of the six different hyperelastic models. The coefficient of detemination measures the correlation between the experimental and model results while the NRMSE gives an indication of the model fitting quality in terms of the fitting error. The evaluation index is a metric that shows how the models fit the experimental results relative to each other. The plotted results show that the polynomial (anisotropic) model performs the best in both the avarage R^2^ and NRMSE values. Besides the highest and lowest levels of the mean for the R^2^ and NRMSE, the polynomial (anisotropic) model also yields the smallest deviations and outliers of its predictions away from the mean line in Figures 2 and 3. With the exception of about three outliers, a large proporportion of the predicted results by the polynomial (anisotropic) model fall within the 95% confidence interval bounds.

**Figure 2.**
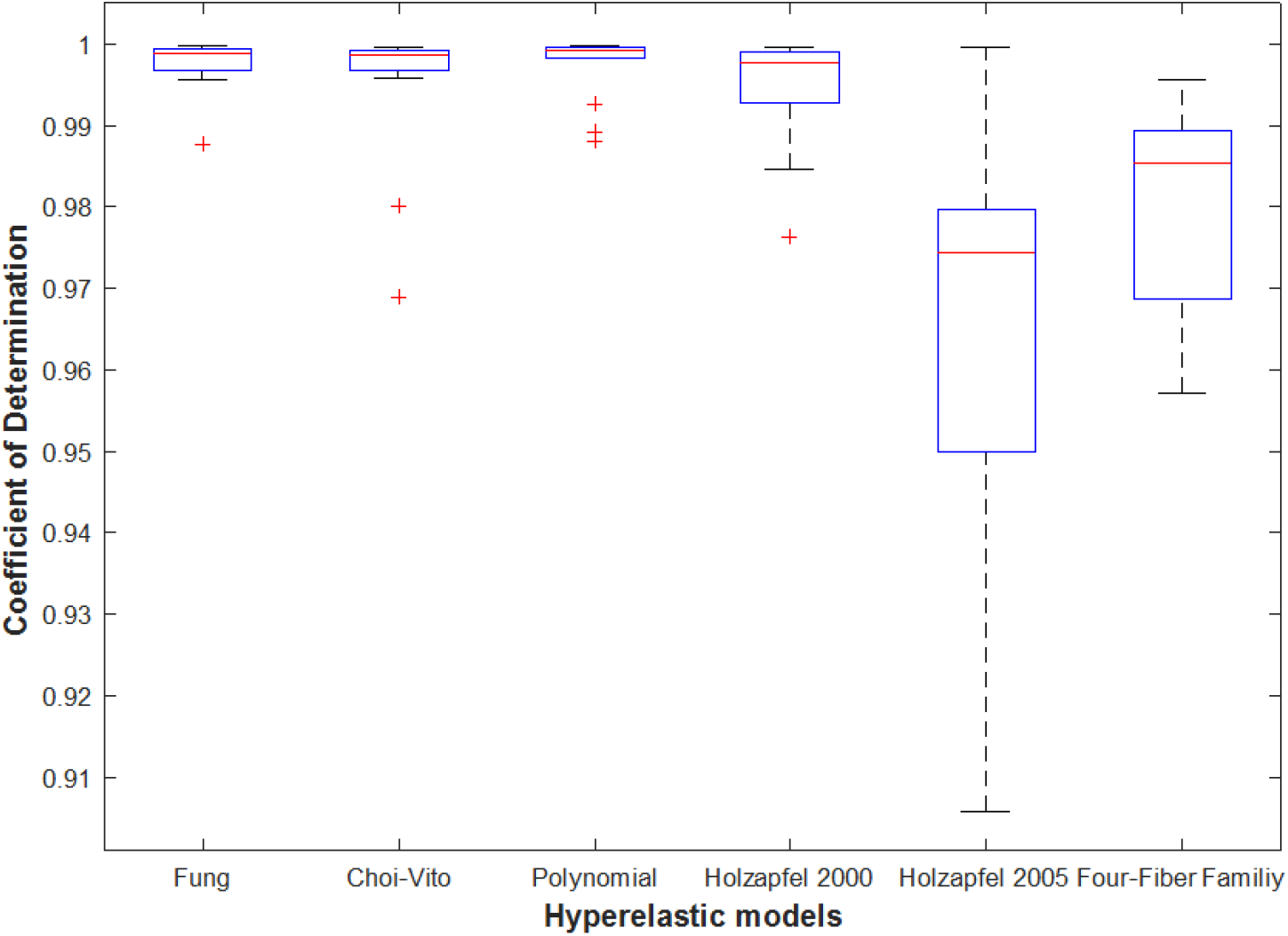
*Average* Coefficient of determination (R^2^) was determined using Equation (9) for N = 13 of the sheep trachea soft tissue for Fung, Choi-Vito, Holzapfel (2000), Holzapfel (2005), Polynomial (Anisotropic) and Four-Fiber Family hyperelastci constitutive models. Holzapfel (2005) hyperelastic constitutive model showing the highest variation of the average R^2^ and Polynomial (Anisotropic) hyperelastic model Showing lowest value of variation.

**Figure 3.**
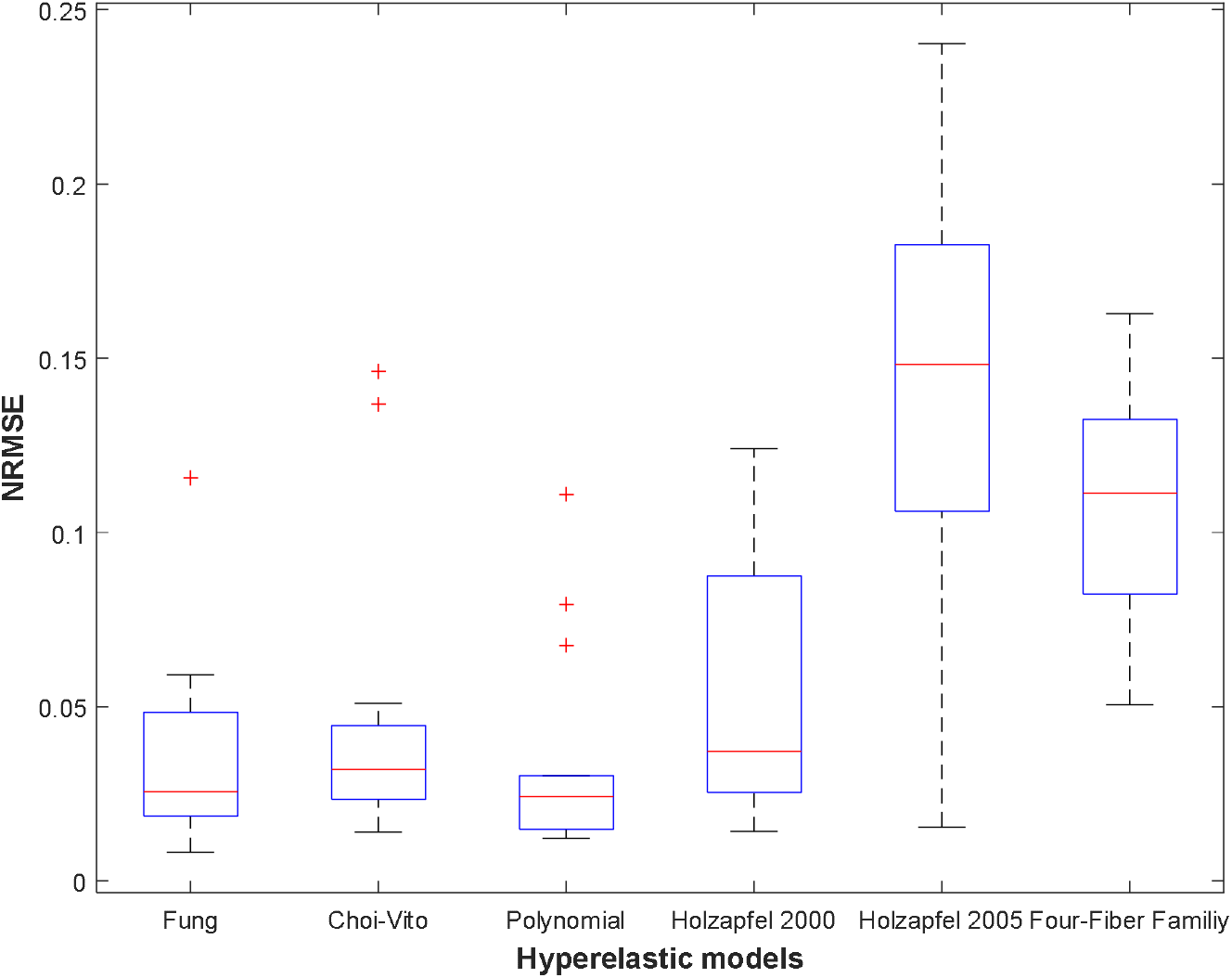
Average Normalised RMS Error (NRMSE)was determined using Equation (13) for N = 13 of the sheep trachea soft tissue for Fung, Choi-Vito, Holzapfel (2000), Holzapfel (2005), Polynomial (Anisotropic) and Four-Fiber Family hyperelastic constitutive models. Holzapfel (2005) hyperelastic constitutive model showing the highest variation of the average R^2^ and Polynomial (Anisotropic) hyperelastic model Showing lowest value of variation.

**Figure 4.**
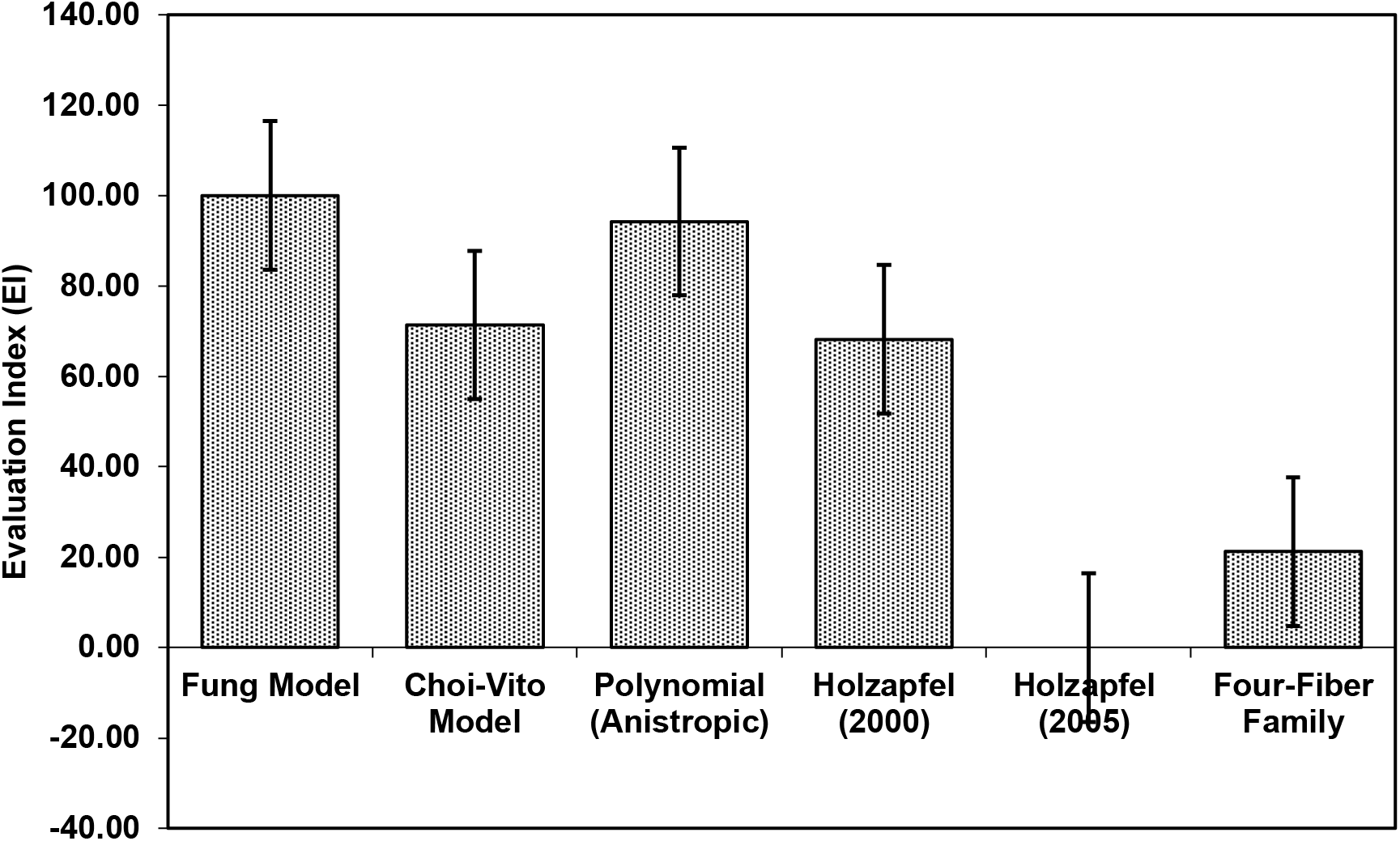
Average Evaluation Index (EI) for all Fung, Choi-Vito, Holzapfel (2000), Holzapfel (2005), Polynomial (Anisotropic) and Four-Fiber Family hyperelastic constitutive models. Holzapfel (2005) hyperelastic constitutive model was found to have the lowest EI as compared to the other five model. Additionally, Fung hyperelastic constitutive model was found to have the highest EI when compared to other models. This means that for this biaxial tensile data, the Fung model has produced the best R^2^.

In Figure 4 the average value of the evaluation index shows that the Fung model yields the best performance (at 100% EI) and that it slightly outperforms the polynomial (anisotropic) model (at around 95% EI). It is interesting to note that both models are implemented as polynomial functions of the material parameters. Of course, the Fung model exhibits a wider range of deviations away from the mean value in their predicted results. The Holzapfel (2005) model yields the worst performance with R^2^ greater than 95%, NRMSE below 20%, and EI equal to zero percent. Since EI is a measure of relative performance of the models, this performance of the Holzapfel (2005) cannot be considered disastrous since it still shows that it is capable of producing correlations above 95% and fitting errors of less than 20%. However, for better accuracy in the predictions of tensile behaviour of the tracheal tissue it might be recommended to apply the Fung or the polynomial (anisotropic) models which yield R^2^ greater than 99.9%, fitting errors NRMSE less than 5%, and EI above 95%. In Figure 4, the Choi-Vito and Holzapfel (2000) have modest performance in the EI at 70% while the four-fiber family yields EI of approximately 20%.

The standard deviations tabulated in Tables 1–6 show that all the material models vary widely from one test to another over the fourteen tests. The amount of standard deviation calculated for each model for each material model is quite substantial. This emphasizes the probabilistic nature of tracheal tissue which shows that it is very hard to apply material parameters from one experiment to predict the tensile behaviour of a different tracheal tissue. As shown in a previous study by the authors, the most appropriate procedure is to use the average values. It is important however to note that these results are only calculated from the circumferential direction of the tracheal tissue.

**Table 1.**
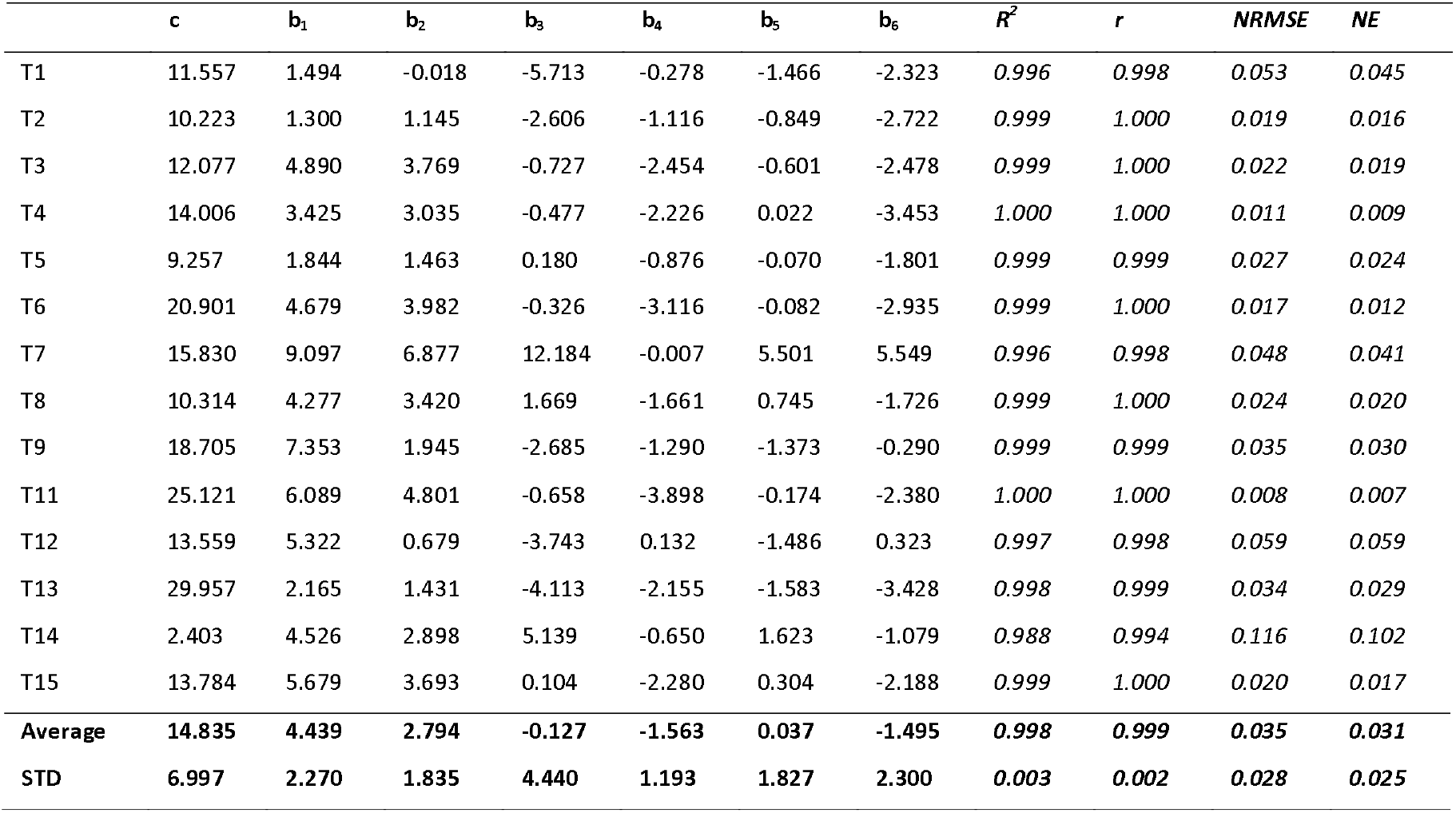
Material parameters of Fung hyperelastic constitutive models (b_1_, b_2_, b_3_, b_4_, b_5_ and b_6_) for diffreent samples of sheep trachea subjected equi-biaxial testing. Coefficient of Determination (R^2^), Correlation Coefficient (r), Normalised Error (NE) and Norm. RMS Error (NRMSE) were also detrmined for all experimental samples.

**Table 2.** Material parameters of Choi-Vito hyperelastic constitutive models (b_o_, b_1_, b_2_ and b_3_) for diffreent samples of sheep trachea subjected equi-biaxial testing. Coefficient of Determination (R^2^), Correlation Coefficient (r), Normalised Error (NE) and Norm. RMS Error (NRMSE) were also detrmined for all experimental samples.

| | $b_0$ | $b_1$ | $b_2$ | $b_3$ | $R^2$ | $r$ | NRMSE | NE |
| --- | --- | --- | --- | --- | --- | --- | --- | --- |
| T1 | 10.871 | 4.730 | 2.299 | -2.159 | 0.997 | 0.999 | 0.045 | 0.039 |
| T 2 | 6.994 | 3.947 | 0.055 | 2.060 | 0.999 | 1.000 | 0.020 | 0.017 |
| T 3 | 5.119 | 10.998 | 1.378 | 6.644 | 0.999 | 1.000 | 0.026 | 0.022 |
| T 4 | 9.512 | 8.210 | -0.216 | 3.711 | 0.999 | 1.000 | 0.022 | 0.017 |
| T5 | 14.711 | 3.732 | 2.111 | -1.334 | 0.998 | 0.999 | 0.037 | 0.028 |
| T6 | 12.812 | 8.204 | 0.059 | 4.001 | 0.999 | 0.999 | 0.027 | 0.023 |
| T7 | 5.229 | 12.525 | 7.776 | -2.304 | 0.969 | 0.990 | 0.137 | 0.113 |
| T 8 | 5.232 | 9.170 | 0.036 | 4.567 | 0.997 | 0.999 | 0.042 | 0.033 |
| T9 | 11.176 | 12.816 | 6.613 | -4.599 | 0.999 | 1.000 | 0.023 | 0.020 |
| T 11 | 13.896 | 8.473 | -0.226 | 3.967 | 1.000 | 1.000 | 0.014 | 0.011 |
| T 12 | 6.335 | 11.095 | 6.644 | -5.919 | 0.998 | 0.999 | 0.037 | 0.031 |
| T13 | 20.311 | 4.876 | -0.239 | 2.209 | 0.996 | 0.999 | 0.051 | 0.045 |
| T 14 | 1.522 | 12.803 | 7.618 | -5.044 | 0.980 | 0.992 | 0.146 | 0.124 |
| T 15 | 6.183 | 12.337 | -0.784 | 5.314 | 0.999 | 1.000 | 0.026 | 0.021 |
| **Average** | **9.279** | **8.851** | **2.366** | **0.794** | **0.995** | **0.998** | **0.047** | **0.039** |
| **STD** | **4.999** | **3.405** | **3.287** | **4.228** | **0.009** | **0.003** | **0.042** | **0.035** |

**Table 3.** Material parameters of Polynomial (Anisotropic) hyperelastic constitutive models (*a_1_, a_2_, a_3_, b_1_, b_2_, b_3_, c_2_, c_3_, c_4_, c_5_, c_6_, φ*) for diffreent samples of sheep trachea subjected equi-biaxial testing. Coefficient of Determination (R^2^), Correlation Coefficient (r), Normalised Error (NE) and Norm. RMS Error (NRMSE) were also detrmined for all experimental samples.

| | $a_1$ | $a_2$ | $a_3$ | $b_1$ | $b_2$ | $b_3$ | $c_2$ | $c_3$ | $c_4$ | $c_5$ | $c_6$ | $\phi$ | $R^2$ | $r$ | NRMSE | NE |
| --- | --- | --- | --- | --- | --- | --- | --- | --- | --- | --- | --- | --- | --- | --- | --- | --- |
| T1 | -1.140 | 1.267 | 0.100 | 3.647 | -0.757 | 1.574 | 1.439 | -0.463 | -0.819 | 4.763 | -1.858 | -0.380 | 1.000 | 1.000 | 0.016 | 0.013 |
| T 2 | 0.992 | 0.873 | 0.634 | 2.634 | 1.325 | 1.923 | 1.439 | 0.353 | 0.414 | 0.277 | -0.065 | 0.264 | 1.000 | 1.000 | 0.012 | 0.008 |
| T 3 | 2.528 | -7.002 | 2.973 | -2.341 | -3.649 | -0.854 | 6.126 | 4.236 | 6.630 | 0.812 | 1.236 | -0.610 | 0.999 | 1.000 | 0.027 | 0.022 |
| T 4 | -0.126 | -4.921 | 1.163 | -0.796 | 5.325 | -2.607 | 9.883 | 3.017 | 5.305 | -5.282 | 6.131 | -0.551 | 0.999 | 1.000 | 0.025 | 0.021 |
| T5 | 1.257 | -1.038 | 0.737 | 3.376 | 0.085 | 0.002 | 3.948 | -2.047 | 1.916 | -0.015 | 0.160 | -0.477 | 0.999 | 1.000 | 0.025 | 0.021 |
| T6 | -6.182 | 17.962 | 1.857 | 5.914 | -4.197 | -4.745 | 21.901 | 9.215 | 0.287 | 5.657 | 2.626 | 0.647 | 0.998 | 0.999 | 0.030 | 0.024 |
| T7 | 2.467 | 18.801 | 6.342 | 8.739 | -0.808 | -6.720 | 8.230 | 6.004 | 2.388 | -1.983 | 7.396 | -0.675 | 0.989 | 0.995 | 0.079 | 0.064 |
| T 8 | 1.536 | 3.701 | 0.307 | -0.581 | 2.517 | -2.001 | 2.554 | 3.278 | 0.344 | -5.450 | 6.187 | -0.286 | 0.993 | 0.997 | 0.068 | 0.058 |
| T9 | 2.091 | 1.905 | 4.896 | -2.114 | 0.633 | -4.655 | 2.877 | 4.360 | 5.348 | 4.407 | -2.354 | -0.307 | 1.000 | 1.000 | 0.012 | 0.011 |
| T 11 | 3.582 | 6.829 | -0.231 | 13.702 | -7.515 | 15.294 | 11.055 | -5.615 | -1.957 | 30.682 | 20.582 | -0.288 | 0.999 | 1.000 | 0.015 | 0.013 |
| T 12 | 0.111 | 2.646 | 3.388 | -0.158 | -0.521 | 1.492 | 0.240 | 4.681 | -4.732 | 5.193 | -0.027 | -0.124 | 0.999 | 1.000 | 0.024 | 0.019 |
| T13 | 2.365 | 3.040 | 0.420 | 10.748 | 3.511 | -6.200 | 3.310 | 10.399 | -1.744 | -1.914 | 0.587 | -0.334 | 1.000 | 1.000 | 0.013 | 0.010 |
| T 14 | 5.177 | -6.558 | 2.761 | -0.703 | -0.407 | 5.269 | -0.482 | 2.674 | -0.678 | 0.993 | 0.797 | 0.797 | 0.988 | 0.995 | 0.111 | 0.093 |
| T 15 | 2.642 | 1.505 | 3.070 | -1.259 | 7.645 | -9.252 | 6.338 | 0.406 | -2.475 | 9.615 | -0.743 | -0.256 | 1.000 | 1.000 | 0.017 | 0.015 |
| **Average** | **1.236** | **2.786** | **2.030** | **2.915** | **0.228** | **-0.820** | **5.633** | **2.893** | **0.731** | **3.411** | **2.904** | **-0.184** | **0.997** | **0.999** | **0.034** | **0.028** |
| **STD** | **2.662** | **7.694** | **1.960** | **5.116** | **3.883** | **6.112** | **5.861** | **4.243** | **3.276** | **8.940** | **5.917** | **0.447** | **0.004** | **0.002** | **0.030** | **0.025** |

**Table 4.**
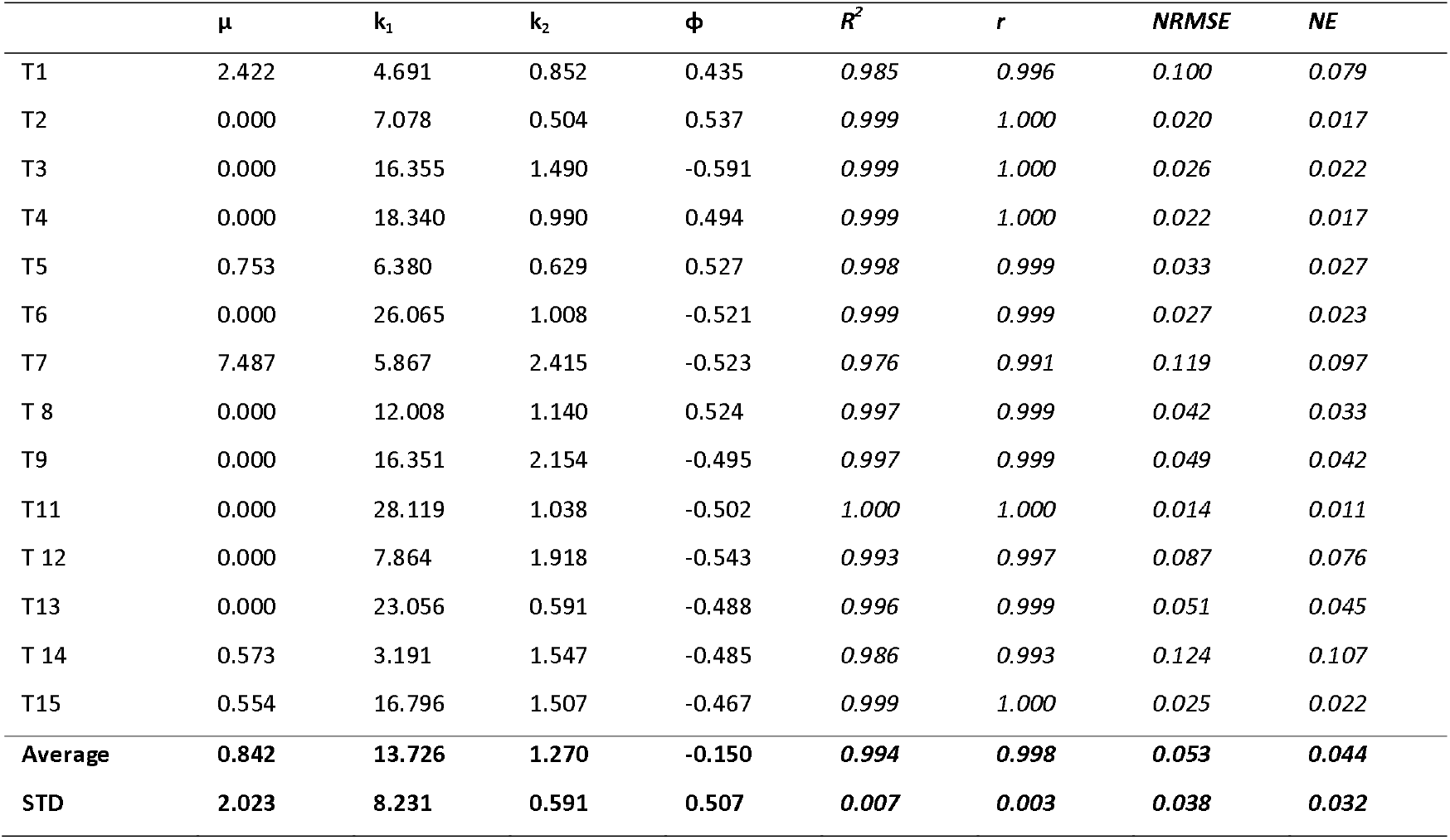
Material parameters of Holzapfel (2000) hyperelastic constitutive models (*μ, k_1_, k_2_ and φ*) for diffreent samples of sheep trachea subjected equi-biaxial testing. Coefficient of Determination (R^2^), Correlation Coefficient (r), Normalised Error (NE) and Norm. RMS Error (NRMSE) were also detrmined for all experimental samples.

**Table 5.** Material parameters of Holzapfel (2006) hyperelastic constitutive models (*μ, k_1_, k_2_, φ, and ρ*) for diffreent samples of sheep trachea subjected equi-biaxial testing. Coefficient of Determination (R^2^), Correlation Coefficient (r), Normalised Error (NE) and Norm. RMS Error (NRMSE) were also detrmined for all experimental samples.

| | $\mu$ | $k_1$ | $k_2$ | phi | $\phi$ | $R^2$ | $r$ | NRMSE | NE |
| --- | --- | --- | --- | --- | --- | --- | --- | --- | --- |
| T1 | 1.992 | 2.040 | 1.186 | 0.253 | 0.895 | 0.967 | 0.986 | 0.149 | 0.123 |
| T2 | 0.242 | 6.636 | 0.422 | -0.462 | 0.906 | 0.999 | 1.000 | 0.021 | 0.018 |
| T3 | 2.773 | 5.590 | 4.342 | -0.657 | 1.351 | 0.978 | 0.996 | 0.106 | 0.092 |
| T4 | 3.748 | 13.618 | 0.943 | -3.141 | 0.758 | 1.000 | 1.000 | 0.016 | 0.013 |
| T5 | 2.439 | 2.489 | 0.840 | 0.056 | 0.758 | 0.981 | 0.995 | 0.105 | 0.087 |
| T6 | 4.123 | 5.919 | 5.421 | 0.664 | 1.595 | 0.940 | 0.985 | 0.185 | 0.154 |
| T7 | 4.153 | 4.317 | 3.067 | 0.526 | 1.028 | 0.976 | 0.993 | 0.120 | 0.100 |
| T8 | 3.246 | 4.610 | 2.994 | -0.559 | 1.205 | 0.973 | 0.994 | 0.132 | 0.117 |
| T9 | 4.934 | 5.947 | 4.619 | 0.530 | 1.235 | 0.953 | 0.989 | 0.195 | 0.160 |
| T11 | 4.459 | 6.191 | 5.865 | -0.619 | 1.574 | 0.932 | 0.985 | 0.181 | 0.156 |
| T12 | 2.347 | 4.430 | 2.852 | 0.542 | 1.077 | 0.978 | 0.993 | 0.152 | 0.129 |
| T13 | 3.211 | 5.432 | 4.423 | 0.661 | 1.547 | 0.906 | 0.972 | 0.240 | 0.191 |
| T14 | 1.773 | 2.145 | 1.446 | -0.121 | 0.799 | 0.980 | 0.991 | 0.147 | 0.129 |
| T15 | 2.951 | 5.491 | 4.619 | -0.588 | 1.394 | 0.950 | 0.988 | 0.183 | 0.158 |
| **Average** | **3.028** | **5.347** | **3.074** | **-0.208** | **1.152** | **0.965** | **0.991** | **0.138** | **0.116** |
| **STD** | **1.241** | **2.820** | **1.847** | **0.991** | **0.306** | **0.026** | **0.007** | **0.063** | **0.052** |

**Table 6.**
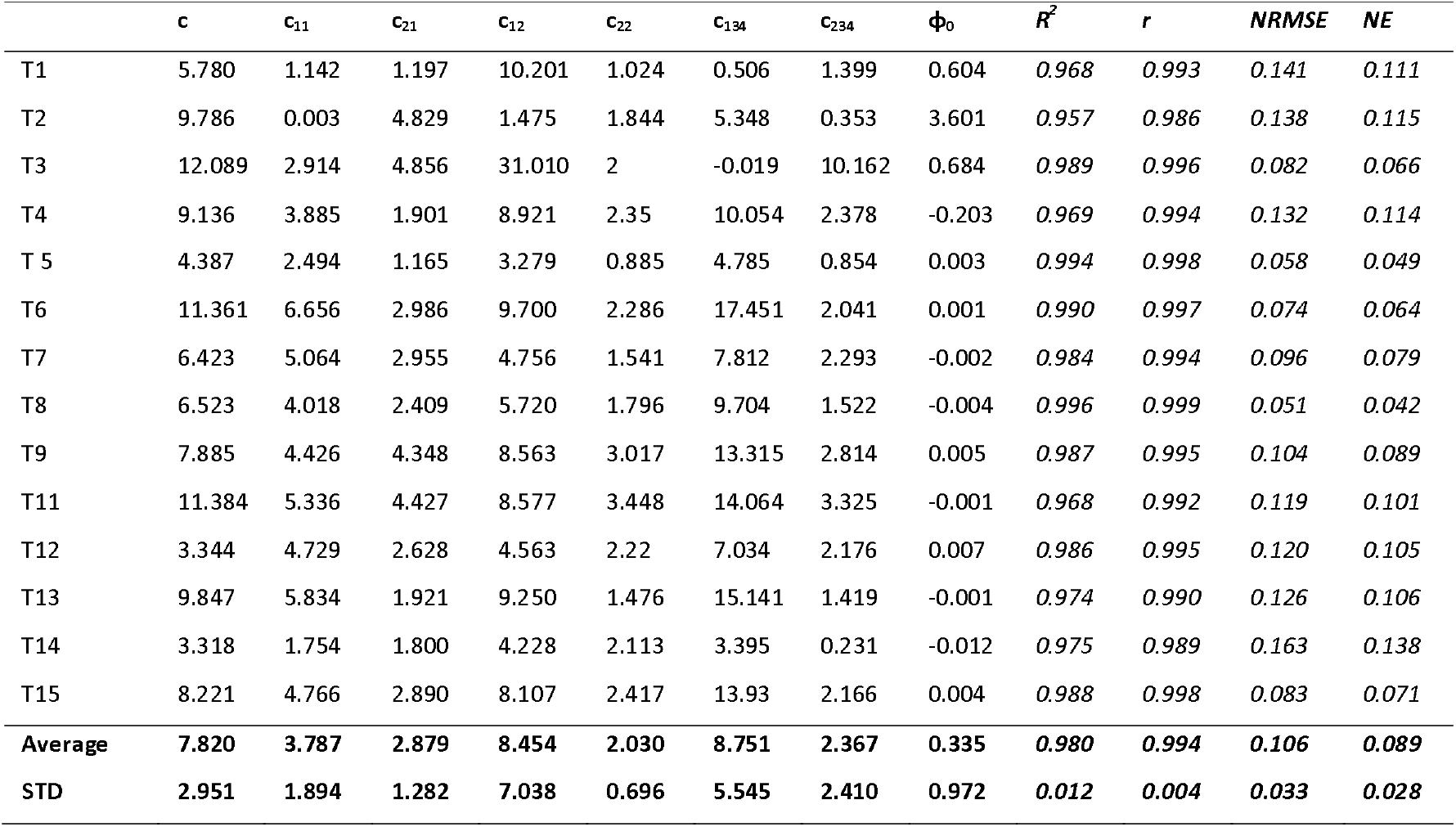
Material parameters of Four-Fiber Family hyperelastic constitutive models (*c, c_11_, c_21_, c_12_, c_22_, c_134_, c_234_ and φ*_0_) for diffreent samples of sheep trachea subjected equi-biaxial testing. Coefficient of Determination (R^2^), Correlation Coefficient (r), Normalised Error (NE) and Norm. RMS Error (NRMSE) were also detrmined for all experimental samples.

## 5. Discussion

The mechanical material properties of the sheep trachea were observed to be nonlinear anisotropic and hyperelastic material, similar to other biological soft tissues [49]. Biaxial testing is a reliable method utilised in understanding the mechanical behaviour of soft tissues [12, 20, 37, 50–52]. The data presented in this study covers the circumferential and longitudinal direction. It is observed that the best performing models are both based on the polynomial implementations having very high numbers of material parameters in their architectures. In our previous studies on soft tissues, the models with exponential implementations such as the two Holzapfel models have yielded much better predictions than the other material models and have always outperformed the Fung model. In this study, the results do not agree with that trend.

Secondly, the performance of the Choi-Vito model with respect to other models within the exponential implementation group is quite remarkable in this study. It performs as well as the Holzapfel (2000) model clearly outperforms the Holzapfel (2005) and the four-fiber family models. Both the Holzapfel (2005) and he four-fiber family models were shown in the oesophagus tissue test to outperform the Choi-Vito model. Thus, for the tracheal tissues, the models that performed poorly in the oesophagus perform much better in the trachea. One of the distinctive features of the trachea is that it consists of rings of cartilaginous tissue embedded within its cylindrical walls. This implies that besides its normal soft tissue which may be dominated by collagen type I, it has other forms of collagen types within its structure. The cartilage tissue is relatively rigid with much less percentage of elongation at fracture. It does not really possess much of the hyperelasticity of the other biological soft tissues. It possesses the property that is desirable for organs that must exhibit some rigidity, elasticity, and lubrication. It is this distinctive property that makes it different from the tensile behaviour of most soft tissues. However, it is not very clear in this study what makes the models such as the Choi-Vito and Fung models perform better than they have in the other soft tissues previously studied by the authors. An understanding of these underlying issues will inform the development of most effective replacement materials for the trachea and all other biological soft tissues.

The performance of the Holzapfel (2005) model over the tracheal tissue is most unreliable. It yields the largest NRMSE value with the widest standard deviation. The polynomial produces the most consistent results and therefore is shown to be the most reliable. However, its debilitating character is that it requires the largest number of material parameters which obviously affects it convergence rates, thereby making the model the slowest and most computationally expensive model of the six material models studied in this research. Therefore, a researcher is more likely to opt for the Choi-Vito or the Fung models with relatively fewer numbers of required material parameters than the polynomial (anisotropic) model.

The higher tissue stiffness along the circumferential direction may be attributed to the effect of the rigid cartilaginous tissue which tends to be pulled axially outwards as opposed to the longitudinal where this tissue has no effect at all. It is therefore likely that tissue stretching along the longitudinal direction was stopped too early. Thus, differential straining may be a better way of testing the trachea than equi-biaxial stretching of the trachea tissue. This is to ensure that there is a fully developed stress-strain curve along the longitudinal direction. It might also be interesting to determine the impact of the cartilage fiber on the total tensile behaviour of the trachea tissue by testing that fiber alone.

The results of the biaxial tension tests of sheep trachea stress-strain response were utilised to fit the strain energy functions of Fung, Choi-Vito, Polynomial (Anisotropic), Holzapfel (2005), Holzapfel (2000) and Four-fiber family model. A direct comparison of hyperelastic constitutive models was made based on evaluation index (EI). The Fung hyperelastic constitutive model was found to have the EI of 100. This means that it is the best performing constitutive model when compared to other five hyperelastic models considered in this study. In addition, Polynomial (anisotropic) hyperelastic constitutive model was found to have the EI of 94 and therefore regarded as second best in terms of performances. This means that both the Fung and Polynomial model were sufficient to capture the nonlinear behaviour of sheep trachea. This was followed by both the Choi-Vito and Holzapfel models (2005, 2000) with EI of 71 and 68, respectively. Finally, the worst performer with EI of 0 was found to be Holzapfel models (2005) model and second last being the Four-Fiber Family with EI of 21.

Despite the contributions offered by this work, some limitations and assumptions should be considered before blindingly applying the results. During the experiment, uniform stress distribution was assumed in the biaxial tests, there will be some stress, especially in the boundary layer.

## 5. Conclusion

This study presents the measured mechanical properties of ex-vivo sheep trachea under biaxial loading. The mechanical data was further utilized for fitting the selected hyperelastic constitutive models. The study has two important findings: firstly, the sheep tracheal tissue is about twice as stiff along its circumferential direction as it is along its longitudinal direction; secondly, the material properties of the sheep tracheal tissue in the different regions are random and different from one another. It would be very interesting if regional dependencies of these material properties were evaluated. Besides the above limitation, the authors believe that the study would have benefitted a lot from correlating these findings with microstructural examinations. Nonetheless, it is believed that the calculated constitutive model parameters will provide more light on the development of accurate computational models for the study of various diseases or medical conditions that affect the trachea. Also, the data presented here will be useful in regenerative medicine and bioprinting of tissue replacements.

## Author contributions

F.N, H.N and T.P contributed to conception, design, data acquisition, analysis, and interpretation, drafted the manuscript, H.N, T.P and F.N contributed to the interpretation of the data and critically revised the manuscript, and finally F.N, H.N and T.P contributed to conception, design, data analysis, and interpretation, drafted and critically revised the manuscript. All authors gave final approval and agree to be accountable for all aspects of the work.

## Additional information

Correspondence and requests for materials should be addressed to F.N. Reprints and permissions information is available at www.nature.com/reprints.

## Acknowledgments

Support from the National Research Foundation (NRF) Grant number (129380) is gratefully acknowledged. Unisa CAPEX Programme supported the acquisition of biaxial testing machine in the Department of Mechanical Engineering, School of Engineering, College of Science Engineering and Technology.

## Corresponding author

Correspondence to Fulufhelo Nemavhola

